# Uniaxial force modifies the length of the mammary ductal network and the orientation of ducts during pubertal development: Findings from computational modeling and laboratory experiments

**DOI:** 10.1101/2024.10.15.618498

**Authors:** Daisy Ulloa, Kelsey M. Teeple, Theresa M. Casey, Uduak Z George

**Affiliations:** Department of Mathematics and Statistics, San Diego State University, San Diego, California, USA; Computational Science Research Center, San Diego State University, San Diego, California, USA; Department of Animal Sciences, Purdue University, West Lafayette, Indiana, USA

## Abstract

The orientation of the epithelium ducts determines the overall shape of the ductal network in the mammary gland, which in turn impacts the efficiency of the delivery of milk through the ducts to breastfeeding infants. However, how the orientations of the ducts are specified is not well understood. Cell-cell and cell-extracellular matrix (ECM) interactions perturb the tissue mechanical environment and influence mechanical signaling, which in turn regulates cell migration, tissue organization, and morphogenesis. This study examined if an applied force that perturbs the tissue mechanical environment can regulate the orientation of the epithelium ducts during puberty, *in vivo*. Uniaxial forces were applied continuously to the right abdominal number four mouse mammary glands (n=10; TEN) at 5-7 weeks of age, that is, during pubertal formation of the epithelial ductal network. The uniaxial force was applied by pulling and adhering the skin around the nipple of the right abdominal number four mammary gland. This pulling force on the right gland made the left abdominal number four gland also experience a contralateral (CONTRA) force. Mammary glands from litter mates were dissected at 3 weeks, 5 weeks, and 7 weeks of unmanipulated mice to serve as controls. Following dissection, whole mounts were prepared by carmine alum staining and panoramic images were captured under light microscopy. The ductal network in the images were skeletonized and straighten along the longitudinal midline to accurately capture the length of the ductal network without bias from its curvature. Findings from using the Measure tool in ImageJ version 1.54j indicated that the length of the ductal network was increased in the TEN and CONTRA mammary glands compared to control (*P <* 0.05). Analysis of the images using OrientationJ version 16.01.2018, an ImageJ plug-in, indicated that the orientation of ducts in the TEN and CONTRA mammary glands where altered compared to control (*P <* 0.05). Although the ductal network was longer in the TEN and CONTRA glands compared to control, there was no significant difference in the total cross-sectional area. In-silico simulations of the ductal network formation using a branching and annihilating random walk model predicted that the increased length of the ductal network may have resulted from changes in the orientation of the epithelium ducts. Thus it is likely that mechanical forces regulate the orientation of ductal branches *in vivo*.

## 1 Introduction

Tissue mechanical environment plays a major role in the proper formation of epithelium and glandular tissues with tree-like structures, such as the lung airways, the salivary gland, pancreas ducts, and the mammary epithelium ducts [1]. These tree-like structures are critical for the long-term physiological functioning of organs and developmental defects often lead to poor health outcomes [2–6]. A great portion of the formation of the lung airways, the salivary gland, and the pancreas ductal network occurs during embryonic and fetal stages of development. Though the formation of the mouse mammary ductal network begins during the embryonic stage of development, it remains rudimentary and composed of a primary duct and 15-20 ductal branches until puberty [7]. The female mammary gland is a great model for studying branching morphogenesis because the expansion from a rudimentary structure to a complex epithelium ductal network occurs postnatally during puberty, allowing for investigations into the evolution of these complex network structures after birth, *in vivo*. The expansion of the mammary ductal network begins at about 4 weeks old in mice, under the influence of hormones. At 10 weeks, the mammary gland is largely infiltrated by a complex network of ducts, which will undergo cycles of branching and regression throughout each estrous cycle [7].

Although it is known that mechanical forces affect branching morphogenesis, a process that governs the formation of tree-like branching structures of ducts within tissues, its effect is not fully understood nor well characterized in the mammary gland. Therefore, how the orientation of the epithelium ductal branches are specified remains an open question [1,8–10]. A crucial step for elucidating how the orientation of the epithelium ductal branches are specified is to identify the mechanisms that modulate or regulate the branch orientation *in vivo*. This would facilitate the selection of potential regulatory candidates and enable the formulation of hypotheses that can be tested through laboratory experiments and computational modeling. Therefore, in this study, we perturbed the mechanical environment and examined how it modifies the orientation of the epithelium ducts in the mammary gland, *in vivo*.

During puberty development of the mammary gland, an extensive network of epithelium ducts emerges from a rudimentary ductal structure via the process of branching morphogenesis (Fig. 1A, [11, 12]). In the mammary gland, branching morphogenesis occurs via two main forms [13]: the first is through successive rounds of elongation and splitting of the tips of existing parent ducts, known as tip bifurcation. The second form of branching is through the elongation of buds that protrude along the sides of existing ducts, known as side branching [14]. Majority of the studies on mammary ductal network formation have focused on identifying the molecular factors that regulate the development of the ductal network including ductal elongation, tip bifurcation, and side branching [15–24]. And comparatively fewer studies aim to elucidate the role of mechanical forces in the formation of the mammary ductal network.

**Fig 1.**
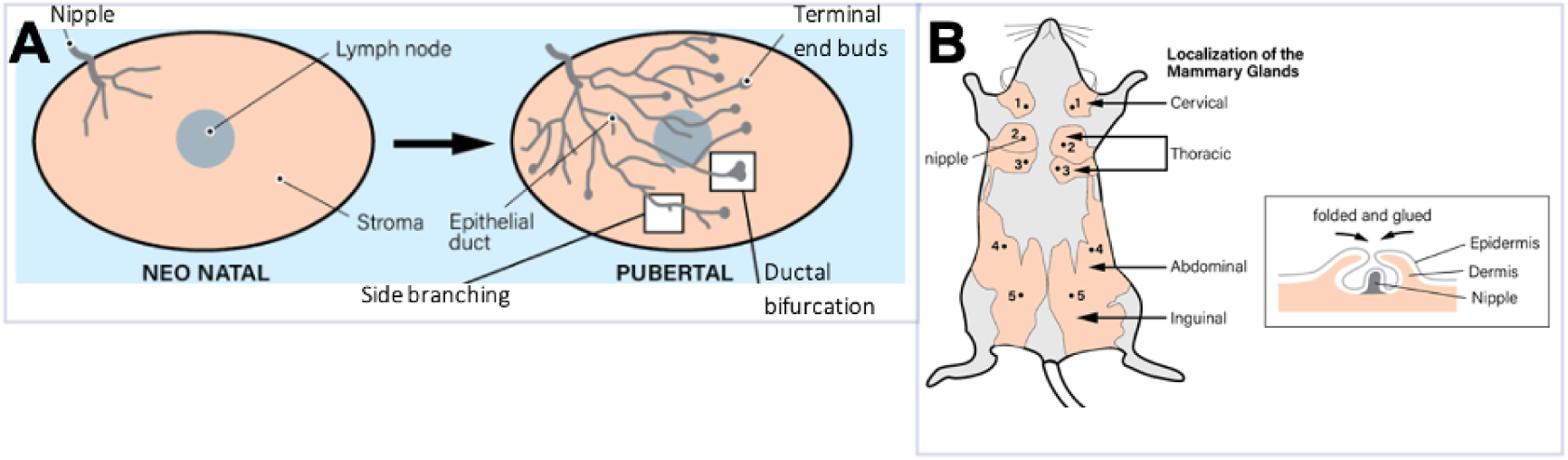
Overview of pubertal branching morphogenesis and laboratory experimentation. (A) Neonatal versus pubertal mammary gland. The formation of an extensive network of epithelium ducts occurs during pubertal development via ductal elongation, tip bifurcation and side branching. (B) The right abdominal number four mammary gland was used to investigate the role of tensional force on mouse branching morphogenesis. The inset shows how a uniaxial tension force was applied to the right abdominal number four mammary gland. The skin surrounding the nipple was lifted, folded and glued together as shown.

Like the developmental changes in the epithelial compartment of the mammary gland, which occur in stages connected to sexual development and reproduction, parallel changes occur in the stromal and fat pad components of the gland [25]. The stroma is relatively acellular, being composed of extracellular matrix (ECM) proteins including fibrillar collagen I, collagen IV and laminins, which play a significant role in controlling cell signaling and the branching process [26–28]. Traction forces exerted by ductal cells reorients and realigns collagen fibers in their preferred direction of migration or tension, facilitating cell movement through the ECM [29]. The reorganization of collagen fibers can in turn alter the stiffness of the ECM, which can affect cell behavior. This creates a feedback loop where cell-generated forces remodel the ECM, which in turn influences cell behavior.

Exogeneous forces such as an applied uniaxial force are also capable of modifying the organization of the ECM. de Jonge et al [30] studied collagen reorganization in fibrin based engineered tissues under uniaxial strain applied continuously for 12 days. By analyzing time-lapse images of collagen orientation, they found that collagen orientation changed gradually from random orientation at day 0 to being parallel to the direction of the uniaxial strain by day 4. Collagen fibers alignment can significantly influence the overall stiffness of the stroma, when collagen fibers are aligned parallel to the direction of an applied load or stress, the tissue exhibits increased stiffness and tensile strength along that direction. Conversely, when collagen fibers are randomly oriented or misaligned, the tissue may exhibit lower stiffness and lower mechanical integrity. Cells can sense the stiffness of the stroma and respond through the process of mechanotransduction. Mechanical signaling can lead to functional and structural responses in the cells and tissues by influencing cell adhesion, migration, morphology and differentiation [31, 32], which are key processes involved in branching morphogenesis [26].

Observations of branching morphogenesis in organoid models demonstrate that the mechanical properties of the region around the ductal branch can directs branch elongation [29] and bias ductal growth along the long axis of the developing gland [33]. Though organoids provide excellent models for studying branching morphogenesis, it is not able to model the complexity of the mammary branched structure and the *in vivo* mammary gland environment [34, 35], therefore *in vivo* studies are needed to further our understanding of the role of mechanical forces in the formation of the ductal network. Therefore, in this study, we aimed to determine whether mechanical forces, applied *in vivo*, can regulate the orientation of the epithelium ductal branches. In order to determine whether mechanical forces can regulate the orientation of the ductal branches, a uniaxial force was applied to the right abdominal number four mammary gland in mice for two-weeks during pubertal development (Fig. 1B, inset). Thereby altering the mechanical environment of the entire gland, including the stroma and fat pad that surrounds the ductal epithelium (Fig. 1A, [36–38]).

Microscopy images of mouse mammary glands subjected to uniaxial force in situ during peripubertal ductal morphogenesis were analyzed and compared to unmanipulated glands to evaluate the effect of the uniaxial force on the ductal network formation. Uniaxial force applied to the right abdominal number four mammary gland induced a contralateral force on the left abdominal number four mammary gland, resulting in unilateral pulling of the left gland, whereas the right gland experienced bilaterally directed pulling. The morphology of the ductal network including the orientation of ductal branches, the length and width of the overall ductal network, the cross-sectional area of the epithelium network, the global curvature of the ductal network and the number of perimetral branches were examined in the glands that experience uniaxial and contralateral forces and were compared to controls. Significant differences occurred in the orientation of the ductal branches, the length ductal network, and the global curvature of the ductal network in the glands exposed to uniaxial and contralateral forces, compared to controls. Furthermore, a branching and annihilating random walk model was applied to study why the ductal network in the mammary glands exposed to uniaxial and contralateral forces where longer than those in controls. Findings from the branching and annihilating random walk simulations indicate that a uniaxial force may regulate the length of the ductal network by modifying the orientation of the ductal branches. This study contributes to increasing our understanding of the effect of mechanical forces on the formation of the epithelium ductal networks.

## 2 Materials and methods

### 2.1 Laboratory experimentation: Application of the uniaxial tension force to the mammary glands

Prior to initiating this study, the protocols were reviewed and approved by Purdue University’s Institutional Animal Care and Use Committee (protocol #190100845). Mice (n=32) used in this study were offspring born to Wap-cre (B6129-Tg(Wap-cre)11738Mam/J) and wild type C57BL/6J crosses (Jackson Laboratory, Bar Harbor, ME) bred in the Casey laboratory at Purdue University. Genotypes of offspring produced by this cross are either wild type or heterozygous Wap-cre. Heterozygous Wap-cre offspring do not show an overt phenotype and animals were not genotyped for this study. Female offspring were euthanized via CO_2_ inhalation at 3 weeks of age (n=6) and at 5 weeks of age (n=6) and their mammary glands were inspected visually in order to confirm that the mammary glands were undergoing normal development and the stage of development was consistent with the age of the mice. The remaining female offspring (n=20) were randomly assigned to one of two groups: control (CTL, n=10) and experimental group (n=10). At five weeks of age, the female mice were anesthetized using 3% isoflurane gas with a flow rate of 1.0 L/minute oxygen. The fur surrounding the nipple of the right abdominal number four mammary gland was removed by plucking with a forcep from all mice (both CTL and experimental groups). The skin on either side of the right nipple was lifted and glued together using topical tissue adhesive (GLUture, Zoetis In., Kalamazoo, MI, USA) to encase the nipple (Fig. 1B), exerting a uniaxial force on the mammary gland. These glands were referred to as TEN glands (n=10); the left abdominal number four mammary gland was referred to as the contralateral gland (CONTRA; n=10). The glue was checked daily. If glue was removed, the mice was anesthetized and skin was lifted and glued again. At 7 weeks of age, the mice in the experimental and control groups were euthanized and their mammary glands examined to determine the effect of the applied uniaxial (TEN) and contralateral (CONTRA) forces on the orientation and distribution of the epithelium ducts.

### 2.2 Collection of the mammary glands and whole mount carmine alum staining

Immediately after euthanasia, the right abdominal number four mammary glands were removed in both the CTL and TEN groups. Although the left abdominal four mammary glands in the TEN groups were not directly exposed to tensional forces, the gluing procedure pulls laterally on the skin beside its nipple thus indirectly exposing these glands to mechanical stress via contralateral forces. Therefore, the left abdominal four mammary glands were also extracted from the experimental mice for analysis and quantification of the effect of the contralateral forces on mammary gland branching morphogenesis. The extracted glands were placed on a plain frosted microscope slide (Fisher Scientific, Hampton, NJ, USA) and gently spread out with the blunt end of the forceps. After allowing the mammary whole mount to dry and adhere to the microscope slide for about 5 minutes, the slides were placed in Carnoy’s solution (75% absolute ethanol and 25% glacial acetic acid) to fix overnight. The next day, the slides were transferred into carmine alum (Stemcell Technologies Inc., Vancouver, British Columbia, CA) and stained overnight at room temperature. The following day, slides were moved into destaining solution (70% absolute ethanol, 5.64% 37% HCl, and double distilled water) for 2 h. Next, slides were dehydrated in increasing concentrations of ethanol (70%, 80%, 95%, and 100%) for 30 minutes at a time. Lastly, slides were defatted in toluene for 30 min or until the fat was sufficiently cleared from the glands. A cover slip was applied using DPX mounting media (Sigma-Aldrich, Burlington, MA, USA) and dried overnight. Once the whole mount carmine alum slides finished drying, slides were viewed with a M60 stereo microscope with a camera port (Leica Microsystems, Wetzlar, Germany). Images from the microscope were captured using a C-mount AmScope camera and saved using the AmScope software (AmScope, Irvine, CA, USA). Multiple images were taken to capture the entire gland by moving the slide across the span of the tissue. To create a single panorama image of the entire gland, Adobe Photoshop (V 23.0.1) was used to merge the individual images into a single image. The microscopy images of the entire mammary gland were used for morphometric analysis for CTL, TEN and CONTRA glands.

The mouse mammary gland has a fairly negligible depth when compared to its length and width and is often considered to be a planar tissue. Therefore, studies in the literature on mouse mammary gland morphometry are often done on planar images of the gland because the planar images provide adequate information without the additional cost that is usually associated with capturing volumetric images of the gland. A volumetric image refers to a visualization of a series of stacked two-dimensional (2D) images that allow for a volumetric representation of a three-dimensional (3D) object. Moreover, except for the area of the ducts, the other key morphometrics (i.e. ductal branch angles, ductal network length) used in this study are not expected to change in value albeit significantly if computed in 3D versus 2D.

### 2.3 Morphometric analysis of the ductal network in TEN, CONTRA and CTL glands

Morphometrics such as the area, length and global curvature of ductal networks and the orientation of the epithelium ducts in the ductal network were computed (Fig. 2).

**Fig 2.**
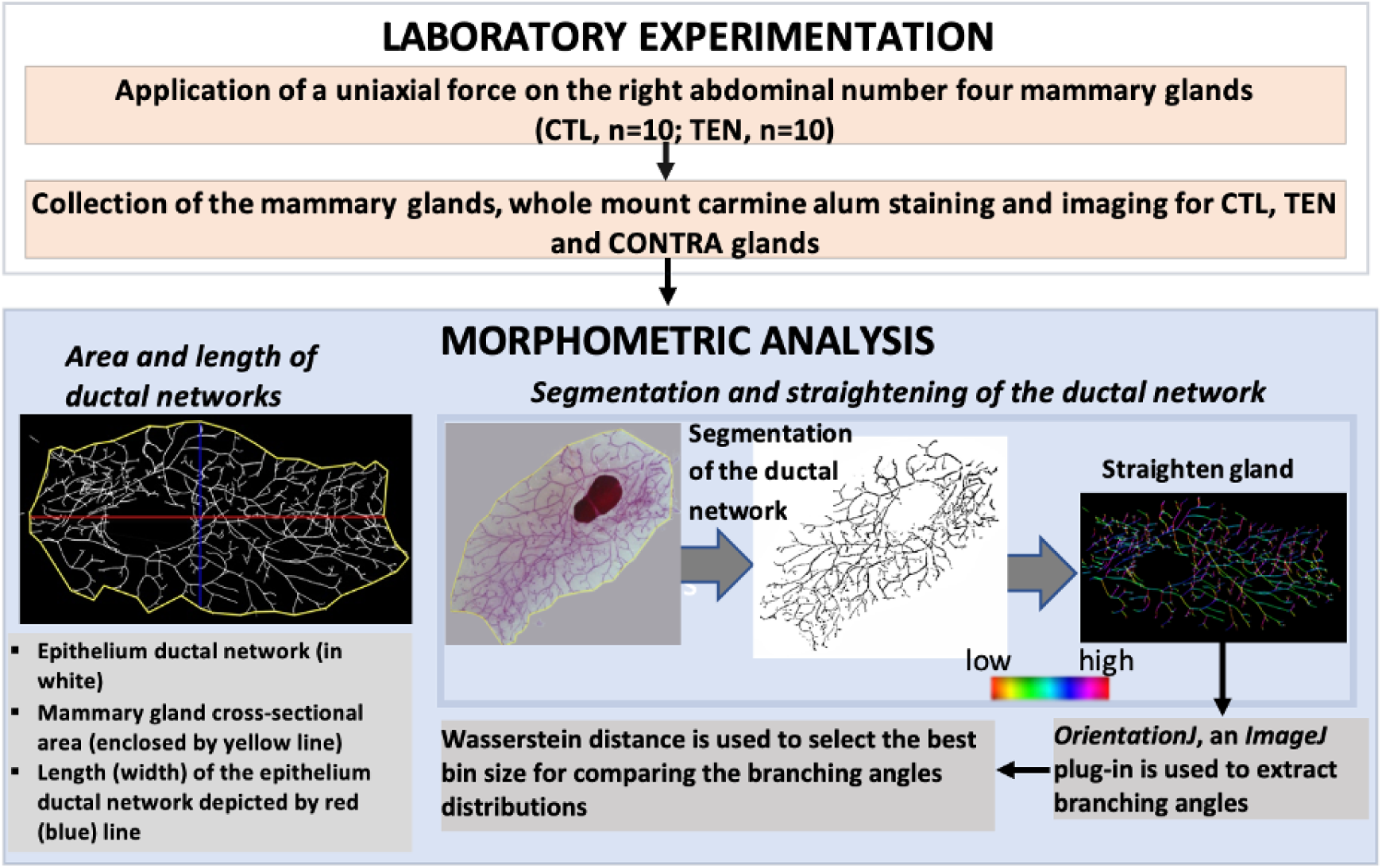
Laboratory experimentation and morphometric analysis. The color map shows high and low branching angles. Morphometrics was obtained for all of the CTL, TEN and CONTRA glands.

#### Global curvature of the ductal network

The images of the entire mammary gland were used to compute the global curvature (also referred to as average curvature) of the ductal network in CTL, TEN and CONTRA glands. This was done using the Kappa Plugin in Fiji open-source image processing package [39, 40]. The average curvature was computed as follows: points where placed on the mammary gland images to trace the longitudinal midline, using the Kappa Plugin, the average curvature was computed by averaging the curvature of each point.

#### Area and length of the ductal network

The total cross-sectional area of the ductal network for each gland was obtained using ImageJ version 1.54j [41] by thresholding the microscopy images to separate the epithelium ducts from the stroma (Fig. 2). After this was completed, the total pixel in the segmented epithelium ductal network was computed by using the Analyze Particles tool in ImageJ to determine the total cross-sectional area. A similar approach was used to calculate the total crosssectional area of the whole mammary gland. Length of the long axis of the ductal network (Fig. 2) were taken using the Measure tool in ImageJ. To accurately compare the length of ductal networks without bias from their global curvature, the branched network were computationally straightened to remove global curvature and ensure that the long axis of the branched network was oriented along the x-axis [33]. To straighten the branched network, we used the straighten tool in the edit menu in ImageJ. After the branched network was straightened, the Measure tool was used to measure the length of the straightened ductal network along the long axis, from the tip of the leftmost branch to the tip of the rightmost branch.

#### Orientation of the mammary ducts

The orientation of the mammary ducts were computed using OrientationJ version 16.01.2018 [42–44], an ImageJ plug-in. In order to compute the orientations of the mammary ducts, the outline of the ductal network was manually traced out in Adobe Photoshop and the resulting skeletal outlines were straightened along the long axis using Fiji’s Kappa plug [39]. Next, the ductal network was adjusted so that the ends of ductal network in all the images pointed in the same direction, for consistency. Maintaining the consistency was important for establishing a common reference for computing and comparing ductal orientations between TEN, CONTRA and CTL glands. Finally, after adjusting the ductal networks to point in the same direction, the orientation of the mammary ducts in the images was computed. This was carried out using the Analysis and Distribution tool in OrientationJ.

In the Analysis and Distribution tool, the variable local window *σ* was set to 10 pixels when computing the local orientations of the mammary ducts. The choice of the value of *σ* can significantly affect the resolution of the orientation analysis. The *σ* value specifies the size of the neighborhood around each pixel that is considered when computing the ductal orientation. *σ* was set to 10 pixels based on work from studies in the literature that demonstrate that this value of *σ* is adequate [42–44]. In OrientationJ, the minimum coherency and energy level was set to 0.1%, meaning pixels were detected if their levels were at or above this percent value. The angle distribution obtained from OrientationJ provided information on the number of pixels that were oriented in each integer degree value between [−90, 90] (Fig. 2). These data provide a sum equal to the total pixels in an image, and the bar graphs generated for data accurately describe the spatial variations in the orientation of the ducts. To find the proportion of branches that lie within a certain degree range, the sum of pixels in the chosen degree range was divided by the total number of pixels in the image. The proportion of ducts in each degree angle was used as a measure for the orientation of the ducts in the TEN, CONTRA, and CTL glands. The distribution of the proportion of ducts in each angle range was compared between TEN, CONTRA and CTL glands. Wasserstein distance, a measure from the optimal transport method, was used to determine the degree of similarity in ductal orientation between TEN, CONTRA, and CTL glands. Wasserstein distance was also used to choose the most suitable number of histogram bins for nuanced similarity comparisons.

### 2.4 Optimal transport method for computing similarities in ductal orientations between TEN, CONTRA and CTL glands

Here we introduce a novel approach that utilizes the optimal transport method to compute the similarities in ductal branch orientations between mammary ductal networks. Optimal transport theory provides a measure to compute distances between pairs of probability distributions. This measure, known as the Wasserstein distance *W* (*µ, η*) satisfy conditions that are desirable when comparing the similarities between two probability distributions, *µ* and *η*. One of the desirable condition is the symmetry condition *W* (*µ, η*) = *W* (*η, µ*) [45]. Another condition that *W* (*µ, η*) satisfies is *W* (*µ, η*) = 0 if *µ* = *η,* which enables the measurement of similarities between two probability distributions. Another commonly used metrics for computing the similarity between probability distributions is the Kullback–Leibler (KL) divergence. The KL divergence *D_KL_*(*µ*||*η*) calculates a score that measures the divergence of the probability distribution *µ* from *η* defined on the same space *ξ*. If *µ* and *η* are discrete probability distributions, then 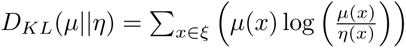 [46]. A drawback of the KL divergence is that it does not satisfy the symmetry condition, that is *D_KL_*(*µ*||*η*) ≠ *D_KL_*(*η*||*µ*). That is, the divergence from distribution *µ* to *η* is not equal to the divergence from *η* to *µ*. Therefore, it is not a good approach for comparing similarities between our probability distributions and is often used as a measure of discrepancy between two distributions rather than a metric [46]. When measuring the similarities between the probability distributions for the branching angles between TEN vs. CTL and CONTRA vs. CTL, the Wasserstein distance was applied because it satisfies the symmetry condition.

Let *µ* represent the probability distribution for the branching angles for the TEN glands and *η* the branching angles for the CTL glands. The probability distribution *µ* (or *η*) is computed by calculating the proportion of branching angles of the TEN (or CTL) glands that lies between different angle ranges. The area under the curve of the probability density function (PDF) for *µ* and *η* are equal and *W* (*µ, η*) computes the optimal cost of moving the contents of *µ* to obtain *η*. The number of bins in *µ* and *η* is represent by *n*. The location of a bin in *µ* is denoted by *x_i_* and the location for all the bins in *µ* as 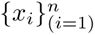. Similarly, location of the bins in *η* is denoted by 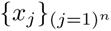. The cost *C_ij_* for moving one unit of item from bin *i* to bin *j* (or vice versa) is measured by the squared Euclidean distance: *C_ij_* = ∥*x_i_* − *x_j_*∥, where 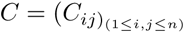 is a symmetric matrix in ℝ*^n^*^×*n*^. A transportation plan *T* ∈ ℝ*^n^*^×*n*^, is defined to describe how many units of item to move from *x_i_* to *x_j_*. The total cost of a given transport plan is:

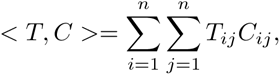

And the optimal transport plan *T* ^∗^ is obtained by solving the following optimization problem:

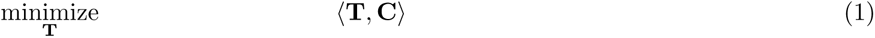

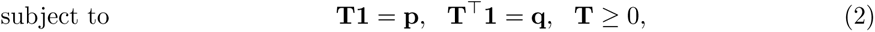

where **1** denotes a vector of ones. The Wasserstein distance 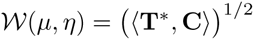, is used to measure the degree of similarity in branching angle distributions between TEN versus CTL glands and CONTRA versus CTL glands.

### 2.5 Modeling ductal network orientation with a biased branching and annihilating random walk model

The biased and annihilating random walk model was used to simulate ductal network formation and study how changes in the branch angles affect the morphology of the ductal network. The most simple random walk moves forward in space, with each step taken in a randomly chosen direction. Random walk models can be described by two attributes: correlated/uncorrelated and biased/unbiased. Correlation refers to whether the direction of each step is influenced on the previous step; bias refers to whether each step has rules that influence direction, while still incorporating randomness in each step [47]. Random walk models can be used to describe movement, such as Brownian motion, as well as cellular and molecular motion [47]. Hannezo et al [48] proposed that correlated/persistent random walks can be used to simulate the branched structure of the ductal network. They used simple rules of branching (i.e. replication) and annihilation (i.e. branch death), to create branching structures similar to those observed in mammary gland experiments. Their model involved three basic rules that defined the dynamics of the tip of the ductal branches : (i) ducts grow outward from active tips, advancing in random directions with a speed *v*, (ii) at any point in time, active tips may randomly split into two branches with a constant probability, (iii) growth ceases for an active tip, when the active tip approaches an existing duct within a defined annihilation distance, causing it to deactivate. Visual inspection of their model simulations of the ductal network revealed good qualitative agreement in spatial organization with experiments [48].

Hannezo et al [48] created an additional random walk that introduced bias toward the distal direction, however, the introduction of bias produced ductal networks that were not in good agreement with experimental data. In a followup study, Nerger et al [33] conducted branching and annihilating random walk simulations that controlled for branching angle between two bifurcating branches. Their simulation results showed that a controlled bifurcation angle is sufficient to reproduce the global bias in the epithelium observed *in vivo*. Here, bias is introduced to tend branching towards the specific angles that we observed to differentiate TEN, CONTRA, and CTL in the laboratory experiments. The goal is to test whether a bias in branching angle alters the length of the ductal network. The rules for the Hannezo et al model include a constant probability for bifurcation and death. Our model does include probabilities for bifurcation and death, however, another rule was included to force a branch to bifurcate if the branch is too long. This was to ensure bifurcations would occur at regular intervals. Similar to Nerger et al [33], we controlled for the bifurcation angle, but at a broader range. Nerger et al controlled for bifurcation angles of approximately 75^°^, whereas we allowed for bifurcation angles from 30^°^ −90^°^ to allow for bias in the movement.

As stated, bias was introduced into the random walk model at each step by limited the randomness of the direction to move towards a given angle. To introduce bias, a function referred to as Biased Random Value, was used. This function works by picking an integer from a window centered about the current angle *α*_0_, and assigning weights to each integer in the window. The window ranges from [*α*_0_ + 15, *α*_0_ - 15] in degrees, where each value in the window is divisible by 5. Values with a higher weight are more likely to be chosen whereas lower weights have less chance of being chosen. The Choices from the Random package in Python version 3.10.9 was used to randomly select a value from the list. The Choices was used because it can apply weights to a list of numbers and then randomly select a number from the list while also taking the weight of each number in to consideration. The biased random value function is described as follows.

#### Biased Random Value function

(i) Set the values for the current branching angle *α*_0_ and the bias *b*_0_. (ii) If say *α*_0_ = 10^°^ and *b*_0_ = 20^°^, then [*α*_0_ + 15, *α*_0_ - 15] := [25,-5] degrees and the window vector, denoting the possible branching angles that the next step in the random walk model can take: is [25, 20, 15, 10, 5, 0, −5] degrees, (iii) reorder values from smallest to largest absolute distance from the bias to obtain [20, 25, 15, 10, 5, 0, −5] degrees, (iv) create list of weights in descending order: [20%, 17%, 17%, 13%, 13%, 10%, 10%] where reordered window list at position *i*, is associated with weight at position *i*, (v) use Choices from the Random package with reordered list and weights as input to choose a new angle value *θ* as the branching angle. Values associated with a higher weight are more likely to be randomly chosen from the list. In this example, angle 20 is more likely to be chosen since the bias is 20.

Our model includes a set of rules for elongation, bifurcation, and intersection events. Each of these rules use a branch head node as input. The head node is defined as the most recent node of an alive branch that the events elongation, bifurcation, or death can occur from. Once an event has concluded, the head node will become a body node that is dormant (Fig. 3A). The rules also depend on information from a matrix that holds the (*x, y*) coordinates of all body and head nodes. This matrix is referred to as Tracker. The Intersection algorithms (Fig 3C) are a series of true/false algorithms that check for intersections between the new head node and existing branches or the boundary during elongation. If the new head node intersects another branch or the boundary, the current head node will die and not elongate to new head node. Otherwise, the current head node will elongate to the new head node.

**Fig 3.**
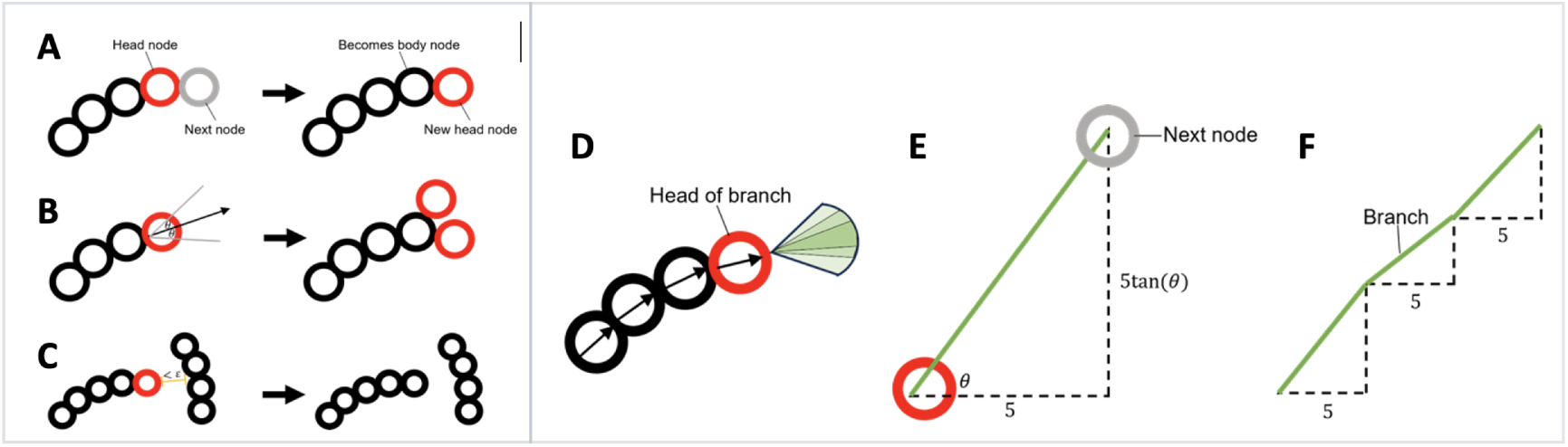
A Biased branching and annihilating random walk model. (A) Shows how head node property changes to a body node as a branch elongates or bifurcates. (B) shows how bifurcation occurs in the algorithm. (C) shows how the intersection functions detect other branches. The concept is the same for intersection of the boundary. (D) shows how the head node will choose the angle for the next node. (E) shows the placement of the next node based on the angle chosen in (D). (F) shows how the branch will form over several elongations

#### Intersection algorithm

(i) retrieve x,y coordinates of head node, (ii) search through Tracker for x,y coordinates near head node, (iii) check if head node intersects with coordinates near head node, (iv) if head node does intersect, return True. Otherwise, return False.

When the bifurcation event is triggered (Fig 3D), two new head nodes will be created from the current head node (Fig 3E). Two new angles are randomized, centered about the angle of the current head node. Then the elongation process is performed starting from the two new angles. The elongation process is described in more detail later.

#### Bifurcation algorithm

(i) use biased Random Value Function to select a value *p*, (ii) Retrieve current angle position of head node *θ* (iii) create two new head nodes with angle position *p* + *θ* and *p* − *θ*, (iii) run elongation algorithm with new head nodes.

The elongation algorithm is how the random walk function is performed. From the current head node, a step of size 5 in the x-direction is taken. The *y*-coordinate is determined by 5 tan(*θ*) where *θ* is the angle of the next step (Fig. 3). The algorithm is defined below.

#### Elongation algorithm

(i) retrieve x,y coordinate (*x*_0_*, y*_0_) and current angle *α*_0_ of the current head node, (ii) use biased random value function with input *α*_0_ to determine new angle *θ*, (iii) new x,y coordinates are defined by (*x*_0_ + 5, 5*tan*(*θ*)), (iv) check if new x,y coordinates intersect another branch or a set boundary by running the Intersection algorithms, (v) if its true the new x,y coordinates intersect, terminate the branch. Otherwise, elongate from current head node to new x,y coordinates. The current head node will become a body node and the new x,y coordinates become a new head node. The new head node is added to the Tracker.

Random walk results were filtered for branching structures that were centered about the horizontal axis, and expressed approximately the same total width. This was done to control for factors that may alter the resulting branching structures, such as curvature and width. This allows for the length of the simulated ductal networks to be compared without bias from the curvature or width of the ductal network.

## 3 Results

### 3.1 Application of uniaxial force did not affect the size of the ductal network and the entire mammary gland

The cross-sectional area of the ductal network and the whole mammary gland was used to estimate their size. The total cross-sectional area of the epithelium ductal network for TEN and CONTRA glands were compared to CTL to determine if the uniaxial force altered the size of the epithelium ductal network. Similarly, the total area of the mammary gland was also compared to determine if uniaxial and contralateral force altered the overall size of the mammary gland. There was no statistically significant difference in the cross-sectional area for the mammary glands in TEN vs. CTL and in CONTRA vs. CTL (Mann-Whitney U-test, *P_T EN/CT L_* = 0.6045*, P_CONT RA/CT L_* = 0.2400, one-tailed). And no significant difference in the area of the epithelium ductal network (Mann Whitney U-test, *P_T EN/CT L_* = 0.6706*, P_CONT RA/CT L_* = 0.3294, one-tailed) (Fig. 4A)

**Fig 4.**
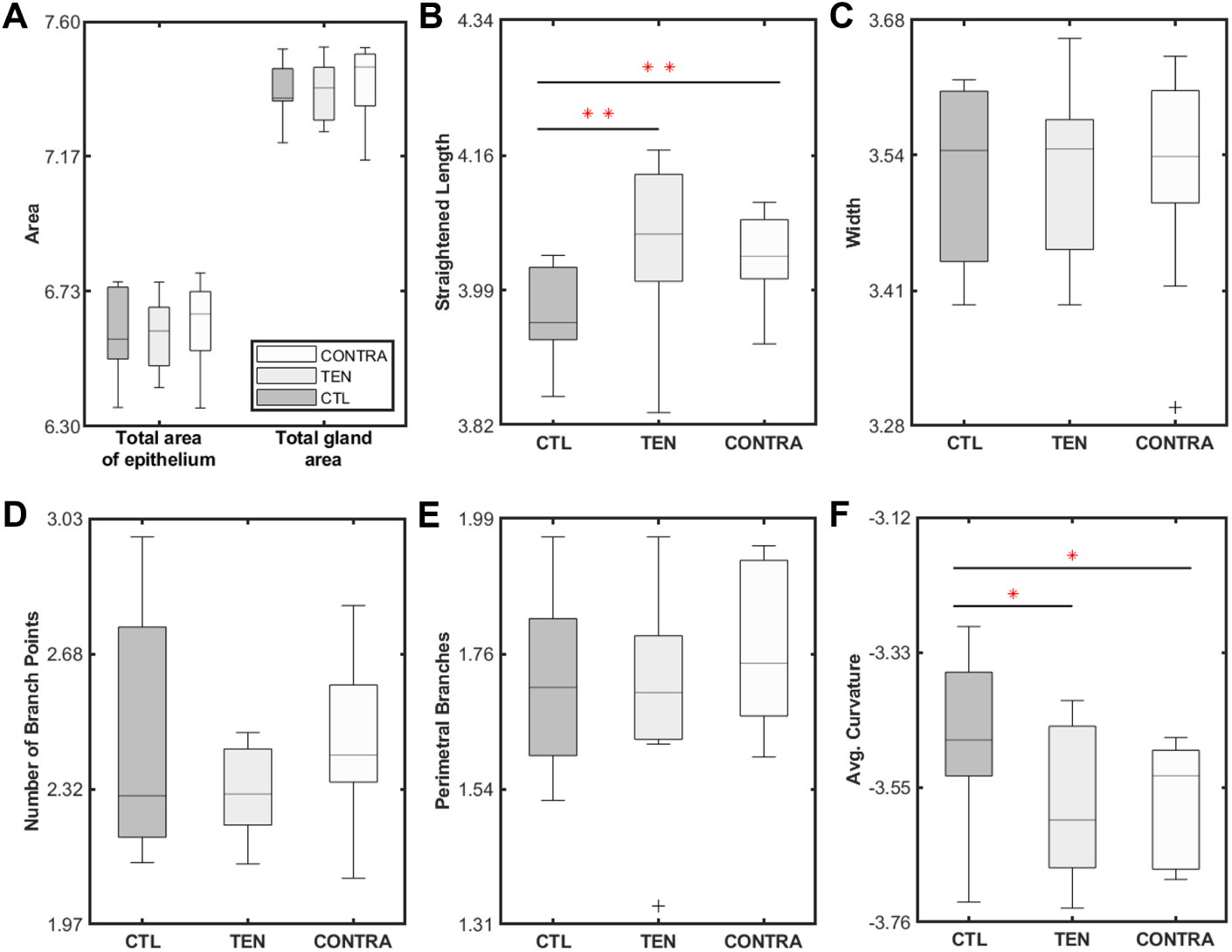
Morphometrics for CTL, TEN and CONTRA glands. All the y-axis is in log base 10. Red asterisk implies that there is a statistical significance difference between the two groups. * represents P *<* 0.1, ** represents P *<* 0.05.

### 3.2 Uniaxial force significantly increased the overall length of the epithelium ductal network for the TEN glands

The length of the epithelium ductal network, measured along the long axis, was significantly longer in TEN and CONTRA compared to CTL glands (Fig. 4B, Mann Whitney U-test, *P_T EN/CT L_* = 0.02*, P_CONT RA/CT L_* = 0.02, one-tailed). Therefore, the increased length for the ductal network in the TEN and CONTRA glands does not correspond to an increase in the cross-sectional area of the epithelium tissue and the total mammary gland. The CONTRA glands had a median length that was longer than CTL but shorter than the TEN glands (Fig. 4B), with the length of the epithelium ductal network varying with the type of force applied. The uniaxial and contralateral forces did not alter the width of the epithelium ductal network, therefore the increased length of the ductal network observed in the TEN and CONTRA glands did not correspond to an increase in the aspect ratio (i.e. the ratio of the length to the width) of the epithelium ductal network.

### 3.3 Uniaxial force minimized the variance in the number of ductal branch points for TEN and CONTRA glands

An important part of the ductal network are the branch bifurcations and side branches. Therefore, the number of bifurcations and side branches in each of the images for TEN, CONTRA and CTL glands were analyzed to determined the effect of the applied uniaxial force on the number of bifurcation and side branches. Application of the uniaxial force resulted in a smaller variance in the total number of branch bifurcations and side branches in TEN and CONTRA glands compared to CTL (Fig. 4D). There was significantly greater heterogeneity (variance) in the number of branches in CTL versus TEN glands (Levene’s test; P=0.0006).

### 3.4 Uniaxial force altered the orientation of the ducts in the TEN and CONTRA glands

The orientation of the individual ducts in the ductal network determines the spatial connectivity and overall shape of the ductal network. The orientation and spatial connectivity of the ducts affects the delivery of milk to suckling neonates. Based on the physics of fluid flow, efficiency in milk delivery through the ducts depends on the diameter of the ducts as well as the orientation of the ducts. Given the importance of ductal orientation, we sought to determine if the angular positions of the ducts were affected by the uniaxial force.

Fig. 5A-B shows the distribution of the ducts using histograms with bins of 60^°^ width. The first bar in the histogram in Fig. 5A represents the ductal regions with angular position that were located between −90^°^ and −30^°^ for all the TEN versus CTL ductal network. The second bar is for the ductal regions with angular position represents the ductal regions with angular position that were located between −30^°^ and 30^°^. Similarly, the third bar is for the ductal regions with angular position located between 30^°^ and 90^°^. The ductal network for TEN had a higher proportion of ductal region located within [−30^°^, 30^°^], whereas CONTRA had a higher proportion of ductal region located within [−30^°^, 90^°^] compared to CTL glands. CTL had a higher proportion of ductal region located within [−90^°^, −30^°^] and [30^°^, 90^°^] compared to those for TEN and a higher proportion in [−90^°^, −30^°^] compared to CONTRA. Reducing the bin size of the histogram allowed us to determine smaller range of angles that contribute more to the observed changes in ductal network orientation between TEN versus CTL and CONTRA versus CTL (see Section **??** and Table **??** in the supplemental text).

**Fig 5.**
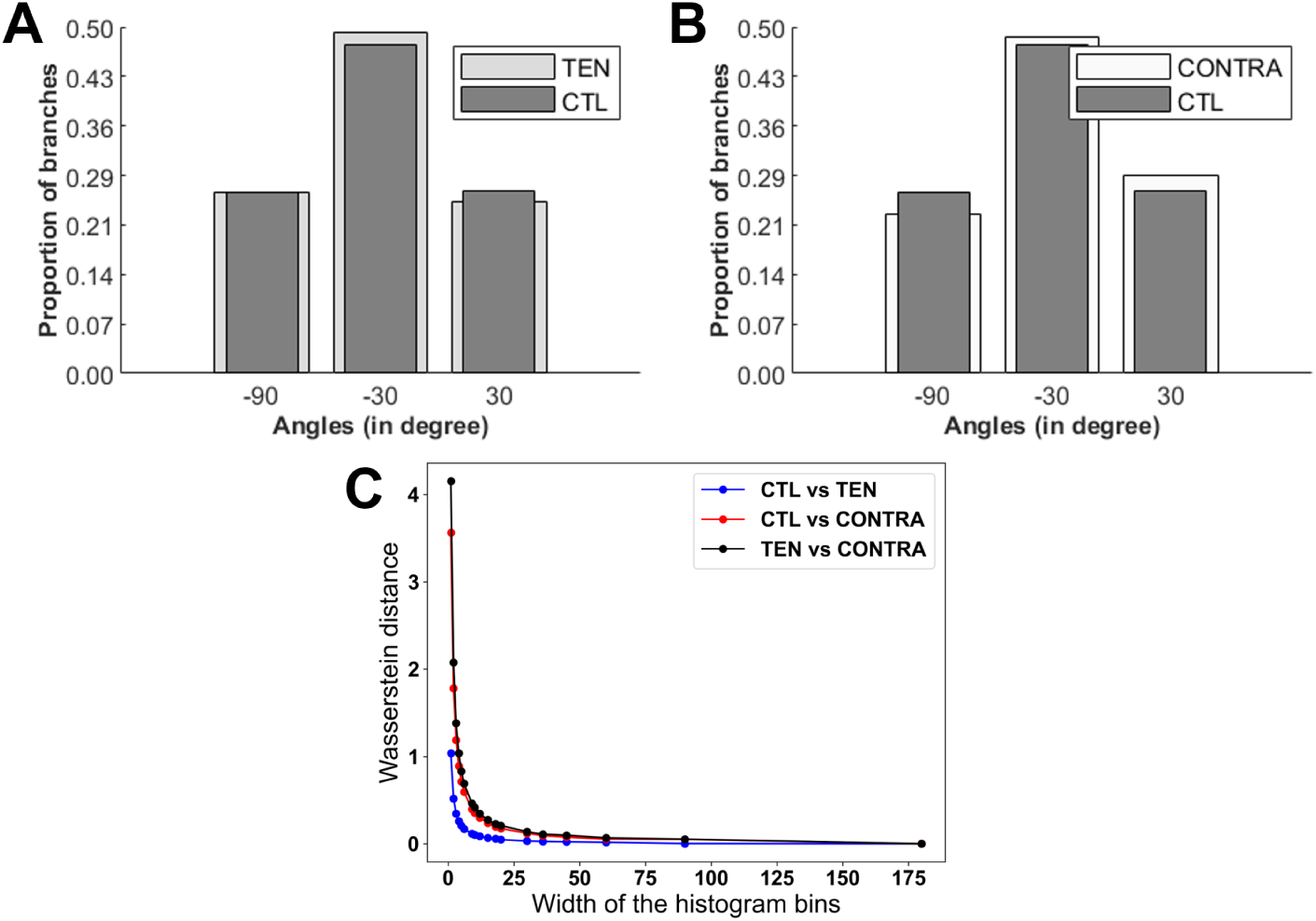
Comparison of the angular distribution of the bulk of the ductal network for TEN, CONTRA and CTL. Visual differences in angle distribution for TEN versus CTL and CONTRA versus CTL are shown in (A) and (B) respectively. Here the bin size is set at 60^°^ angle. (C) The graph of Wasserstein distance against bin size shows how the degree of similarity in angle distribution between the different groups is dependent on the bin size.

Note that if the bin size is very large (say 180^°^), then one will not be able to capture the differences between TEN, CONTRA and CTL ductal network. On the other hand, if the bin size is too small (say 1^°^), one will only learn that each ductal network is different. In order to select a good bin size for comparing the histograms, the similarity in the distribution of the angular positions for TEN vs CTL and CONTRA vs CTL for different bin sizes was computed. The similarity in the distribution of the angular positions for TEN vs CTL and CONTRA vs CTL was measured using a novel approach that utilizes the Wasserstein distance. Wasserstein distance clearly captures the expected dynamics for how the similarity between the angle distributions would change with the histogram’s bin size (Fig. 5C). A low values of Wassertein distance indicates a high similarity between the distributions and a high values of Wasserstein distance indicates a high degree of dissimilarity between the distribution (Fig. 5C). The graph of Wasserstein distance against bin size resembles a visual tool called elbow plot that is often used in cluster analysis and other machine learning algorithms to determine the optimal number of clusters [49]. In particular, elbow plot identifies the point at which adding more clusters does not significantly improve the model performance, indicating a diminishing return. Using a similar idea as often used for elbow plots, a bin size of 5^°^ was selected because it gives a value for Wasserstein distance close to the elbow of the graph in Fig. 5C.

Fig. 6 shows a visual comparison of the angle distribution for the ductal network for TEN versus CTL and CONTRA versus CTL using 5^°^ angles for the bin size. Consistent with the angle distribution with 60^°^ bin size, we see here that TEN glands consistently had higher proportion of ductal regions with smaller angular position (i.e. between [−20^°^, 35^°^], Fig. 6A). A different visual pattern evolve for the angle distribution for CONTRA versus CTL (Fig. 6B). Fig. 6B indicates that the higher proportion of angular regions in [−30^°^, 90^°^] angles was due to a high proportion of ductal regions existing in the [−5^°^, 55^°^] degree range. The CONTRA glands experienced the uniaxial force on one side of its gland, that is on the side adjacent to the TEN gland. The higher proportion of smaller angles was only evident on one-side of the ductal netwok of CONTRA vs CTL (Fig. 6B). Therefore, Fig. 6A-B shows that the uniaxial force altered the orientation of the ducts in the TEN and CONTRA glands.

**Fig 6.**
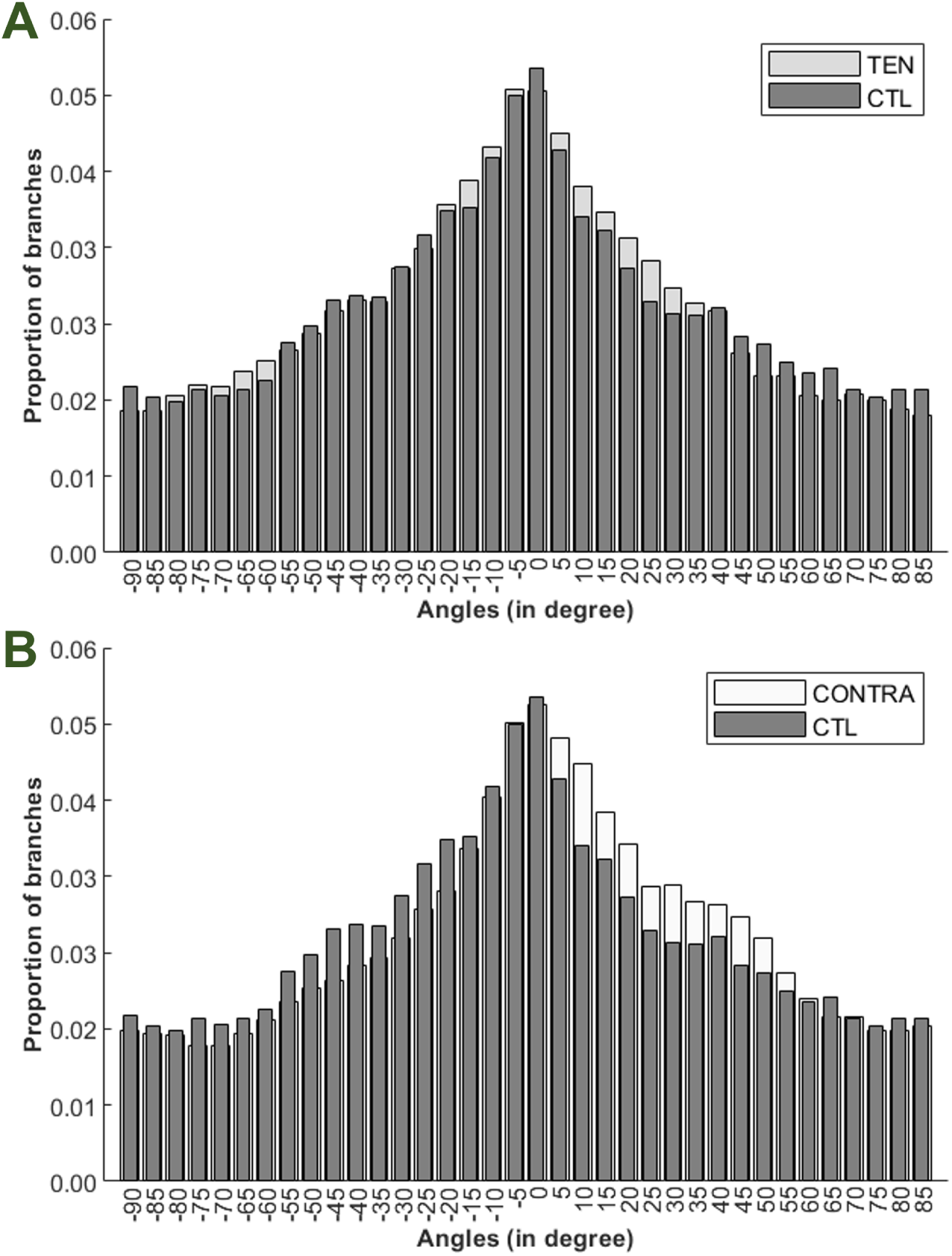
Angle distribution for the whole bulk of the ductal network. Here the bin size is set at 5^°^ angle.

Analysis of the distribution of angles for each individual ductal network, showed that the angle distributions were altered by differing application of forces, TEN and CONTRA, relative to CTL (Fig. 7). The TEN glands, in general, had a significantly higher proportion of ductal branches within the −15^°^ to −11^°^ range of angles. And the CTL glands had a significantly higher proportion of branches within 50^°^ to 59^°^ and 85^°^ to 90^°^ range of angles. CTL had a significantly higher proportion of ductal branching within −75^°^ to −71^°^ and −45^°^ to −31^°^ compared to CONTRA. In contrast, CONTRA had a significantly higher proportion of ductal branching compared to CTL in the following ranges: 10^°^ to 14^°^ and 20^°^ to 29^°^. Thus, the uniaxial force biased ductal orientation differently in CONTRA compared to the TEN glands, indicating that the direction of the applied force is important for controlling the orientation of the ducts.

**Fig 7.**
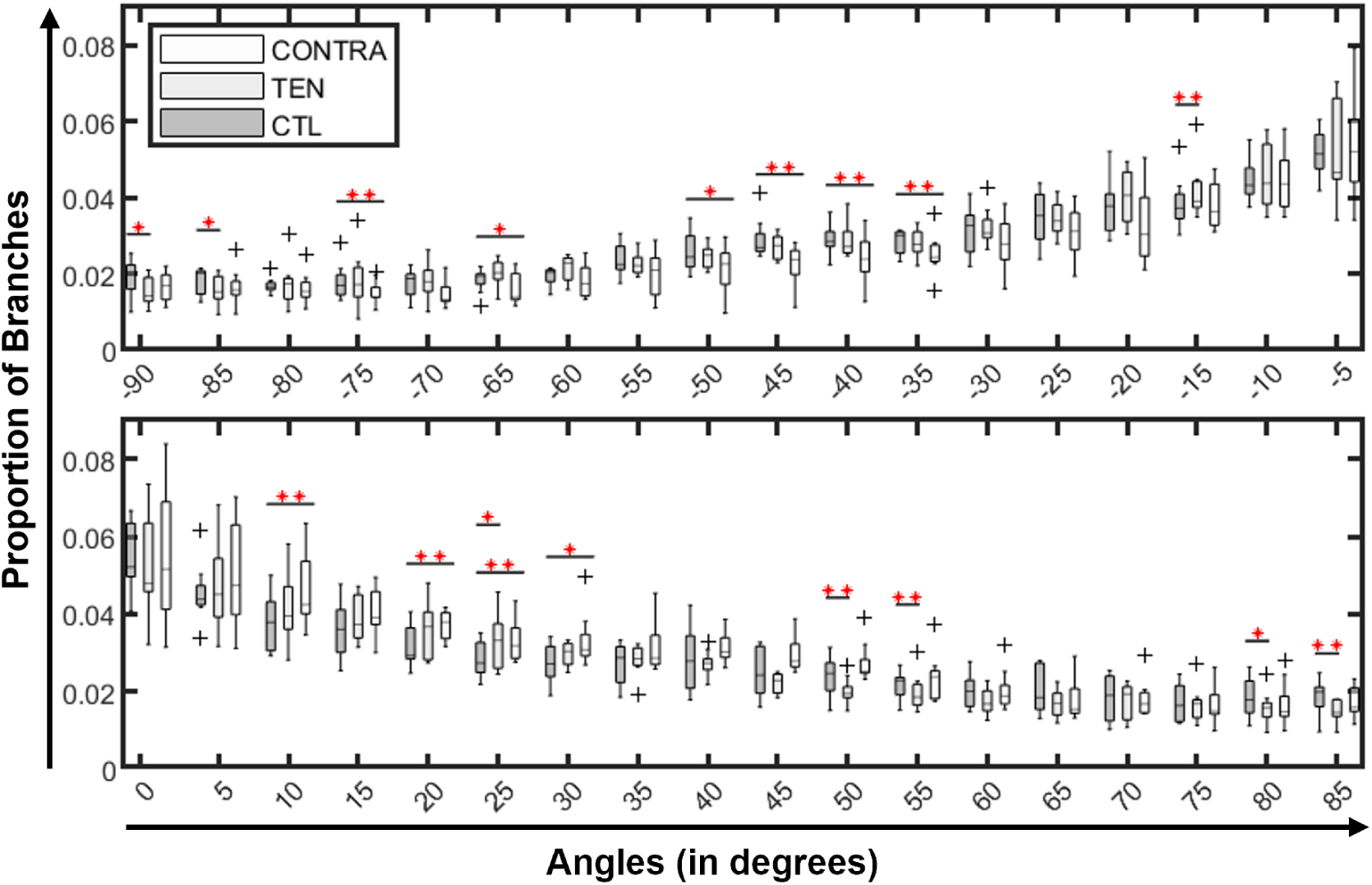
Angle distributions for the individual ductal networks in the TEN versus CTL and CONTRA versus CTL. Here the bin size is set at 5° angle. ** represents P *<* 0.05.

Considering that the orientation of daughter ducts is in general determined by the branching angles of daughter ducts from their parent duct, it was hypothesized that the increased length of the ductal network for the TEN and CONTRA glands may have resulted from changes in the branching angle of the ducts. To test this hypothesis, ductal network formation was simulated using a biased branching and annihilating random walk model and the lengths of the ductal networks were analyzed to determine if varying the angle of branching can alter the length of a ductal network.

### 3.5 Biased and annihilating random walk model predicts that the increased length of the ductal network in the TEN glands was due to the existence of smaller branch angle for ductal bifurcation in TEN compared to CTL gland

The biased and annihilating random walk model was simulated for 18 different instances, 9 for TEN and 9 for CTL. TEN glands were given a bias angle of 15^°^ and −15^°^ whereas CTL was given a bias angle of 50^°^ and −50^°^. The simulations were run for 25 generations of ductal bifurcations, which appeared to be sufficient generations for multiple branch bifurcations and deaths. The length of the simulated ductal networks were computed and compared to those obtained from the laboratory experiments. The results from the simulations predict that smaller angles of ductal bifurcation can significantly increase the length of the ductal network (Fig. 8).

**Fig 8.**
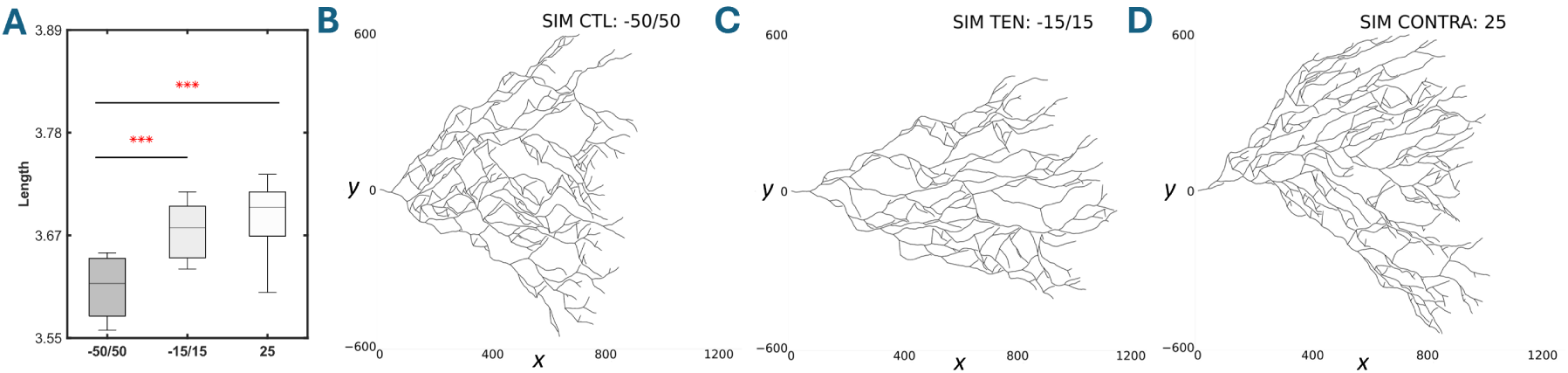
Biased and annihilating random walk simulations. (A) shows the significant difference between the length of the simulated CTL, TEN, and CONTRA glands; y-axis is in log base 10. ( B, C, & D) shows a sample simulated CTL, TEN, and CONTRA gland respectively obtained after 25 generations of ductal bifurcations.*** denotes P*<* 0.005.

Further analysis for one-sided bias was conducted, where the simulation was given only one angle, rather than a positive and negative angle of the same distance from 0^°^. The upper and lower bounds of branching angles between CONTRA and CTL that were significant in the experimental data was used, i.e. −75^°^, −35^°^, 10^°^, and 25^°^. Simulations from the one-sided bias was also compared to the two-sided bias, where a positive and negative angle of the same distance from 0° was used as bias, i.e. using the following pairs of angles −15^°^*/*15^°^, −50^°^*/*50^°^, −85^°^*/*85^°^. Comparisons of the simulations from the one-sided bias with the two-sided bias indicates that small branching angles, one- or two-sided, result in significantly longer ductal networks than ductal networks that prefer to branch at larger branching angles (Table 1). Based on results from 9 different simulations for each of the one-sided and two sided bias. Table 1 also indicates that two branching structures with a preference for small branching angles will not grow significantly longer than each other. The line between small and large branching angle appears to be at or above 35^°^.

**Table 1.** Comparison of the one-sided and two-sided bias simulations of the biased and annihilating random walk model. The values in the Table are p-values from the one-sided Mann-Whitney U test. The p-values indicate the statistical differences in ductal network when testing if the bias angle in row is significantly less than those in the corresponding column.

|  |  | Two-sided bias |  |  | One-sided bias |  |  |  |
| --- | --- | --- | --- | --- | --- | --- | --- | --- |
|  |  | -85/85 | -50/50 | -15/15 | -75 | -35 | 10 | 25 |
| Two-sided bias | -85/85 |  | 0.0927 | 0.0003 | 0.0318 | 0.0040 | 0.0005 | 0.0005 |
|  | -50/50 |  |  | 0.0018 | 0.1657 | 0.0086 | 0.0018 | 0.0010 |
|  | -15/15 |  |  |  | 0.9830 | 0.8343 | 0.2134 | 0.1447 |
| One-sided bias | -75 |  |  |  |  | 0.1447 | 0.0040 | 0.0040 |
|  | -35 |  |  |  |  |  | 0.0789 | 0.0211 |
|  | 10 |  |  |  |  |  |  | 0.2981 |
|  | 25 |  |  |  |  |  |  |  |

## 4 Discussion

Identifying the mechanisms that regulate the orientation of the ductal branches is an important step towards unraveling the complex interactions that coordinate the formation of the mammary ductal network during puberty. A crucial step for elucidating how the orientation of the epithelium ductal branches are specified is to identify the mechanisms that modulate or regulate the branch orientation in vivo. Biomechanical forces originating from cell-cell and cell-ECM interactions actively remodel the tissue mechanical environment, modulating mechanotransduction pathways that regulate cell migration, tissue organization, and morphogenesis. In this study, we investigated if a uniaxial force applied during puberty can regulate the orientation of epithelium ducts in the ductal network. Using a combination of in-vivo experiments and computational modeling, we demonstrated that a uniaxial force applied to mice mammary glands for two-weeks during pubertal development increases the length of the ductal network and modifies the orientation of ducts in the ductal network. The cross-sectional area of the ductal networks that were exposed to the uniaxial and contralateral force (i.e. the TEN and CONTRA glands) were fairly similar to CTL, which made us conclude that the increased length of the ductal network may not have corresponded to a change in the size of the ductal network. In other words, one would expect the cross-sectional area to increase as the ductal network gets longer. Therefore, the applied forces may have changed some measures in the ductal network in order for TEN and CONTRA to have a longer ductal network while having a similar cross-sectional area as CTL.

The orientation and spatial connectivity of the ducts affects the delivery of milk to suckling neonates. Given the importance of ductal orientation, we sought to determine if the angular positions of the ducts were affected by the uniaxial force. The proportion of ducts in each degree angle was used as a measure for the orientation of the ducts in the TEN, CONTRA, and CTL glands. We introduced a novel approach that utilizes Wasserstein distance metrics to identify an optimal bin size for comparing the distributions of ductal angles in TEN and CONTRA glands to CTL. The proportion of ducts at different angular regions in the ductal network for TEN and CONTRA was compared to CTL, and we were able to determine how the orientation of ducts differ in TEN and CONTRA versus CTL. The TEN glands, in general, had a significantly higher proportion of ductal branches within the smaller angles i.e. −15^°^ to −11^°^ range of angles compared to CTL. While CTL glands had higher proportion of branches within the larger angles i.e. within 50^°^ to 59^°^ and 85^°^ to 90^°^ range of angles. Therefore, the uniaxial force in the TEN glands bias the orientation of ducts towards the smaller angle ranges. Which implies that the applied uniaxial force modified the orientation of the ducts during the formation of the ductal network. CTL had a significantly higher proportion of ductal branching within −75^°^ to −71^°^ and −45^°^ to −31^°^ compared to CONTRA. In contrast, CONTRA had a significantly higher proportion of ductal branching compared to CTL in the following ranges: 10^°^ to 14^°^ and 20^°^ to 29^°^. Therefore, the contralateral force in the CONTRA glands also bias the orientation of ducts towards the smaller angle ranges. Thus, the uniaxial force biased ductal orientation differently in CONTRA compared to the TEN glands. This implies that the direction of the applied force influences the orientation of the ducts.

Furthermore, we hypothesized that the increased length in ductal network for the TEN and CONTRA glands may have been due to changes in orientation of the ducts compared to CTL. Since the orientation of daughter ducts is in general associated with the angle of branching of daughter ducts from their parent duct, we tested this hypothesis by examine how variation in the ductal branch angles affect the length of ductal network, using a biased branching and annihilating random walk model. Results from the biased branching and annihilating random walk simulations predict that the increased length of the ductal network in the TEN glands may have been caused by alterations in branching angles. In particular, narrow ductal branch angles led to an increase in the length of the ductal network. Based on our findings, we concluded that the uniaxial force caused an increase in the length of the ductal network of the TEN glands by modifying the orientation of the ductal branches.

Other metrics such as the average curvage and number of ductal branch points were also modified by the application of the forces. The TEN and CONTRA glands had smaller variation in total number of branches within group compared to CTL, which implies that the uniaxial force may have influenced the rate of branching in the epithelium ductal network, though not significantly. Studies on mouse embryonic lung morphogenesis, in which mechanical forces were applied by increasing intraluminal pressure, have shown that mechanical forces can accelerate lung growth and increase by 2-fold the average number of new epithelial branches between 24 and 48 h of culture [50]. Though the total number of epithelium branches in the embryonic lung was doubled, the diameter of the branches were reduced by 30%, and the global size of the lungs remained similar to the control ones [50]. This shows that mechanical forces can change the organization and dimensions of the epithelial branches in the mammary glands and lungs, while the overall size of the epithelial branched network stays unchanged.

This study raises new questions concerning the mechanism by which the uniaxial force regulated the orientation of the ducts. That is, whether the changes in ductal orientation was orchestrated by purely physical interactions between for example collagen fibers and ductal cells or if it was orchestrated by purely molecular factors, a combination of both physical and molecular factors or through entirely different mechanisms. Given that a uniaxial force applied to a tissue can cause a uniaxial strain, resulting in collagen reorganization in the direction of the uniaxial force. And changes in collagen organization can alters mechanical signaling, influence cell adhesion, migration, morphology and differentiation [31, 32], which are key processes involved in branching morphogenesis [26]. Future work may focus on understanding how uniaxial force impact cell behavior during branching morphogenesis. This may yield new insights on how the tension force generated by interactions between the ductal cells and its surrounding ECM affect cell behavior and the process of mammary branching morphogenesis. The findings from this study also highlights the need to further investigate the role of collagen fiber orientation and organization on cell behavior during the morphogenesis of the pubertal ductal network in the mammary gland, because collagen fibers are known to provide contact guidance for cells.

In summary, findings from this study indicates that mechanical forces may regulate the orientation of ductal branches and the overall length of the ductal network. Improving our understanding of the mammary ductal network formation is important because the epithelium ductal network coordinates milk synthesis and secretion in lactating mammals and may also affect long-term milk production, providing the necessary spacing for proliferation of milk producing lobulo-alveolae. The orientation/branching angles of the epithelium ducts determine the overall shape of the epithelium ductal network. An improved understanding of how mechanics impacts the formation of the epithelium ductal network is essential for understanding factors that may lead to developmental abnormalities. It may also yield insights into the formation of other branched organs and may potentially lead to the identification of novel ways to design artificial organs to combat diseases [51].

## 5 Acknowledgments

This project was supported by a National Science Foundation CAREER DMS2240155 grant to U.Z.G. The funders had no role in the study design and implementation, decision to publish, or preparation of the manuscript.

