## Supplemental File for "Uniaxial force modifies the length of the mammary ductal network and the orientation of ducts during pubertal development: Findings from computational modeling and laboratory experiments"

**The PDF file includes:**

Supplementary Text

Tables S1

### Comparisons of the angular positions of the ducts in TEN and CONTRA to CTL for 5°, 10° and 20° bin sizes

In Fig. 5A-B, we showed the distribution of the angular positions of the ducts using histograms with bins of 60° width. To further determine how the bin size affects the ability to detect the differences in the angular positions of the ducts in TEN and CONTRA compared to CTL, we analyzed the differences for bin size less than 60°. Table S.1 shows the angular positions that were significantly different in TEN and CONTRA compared to CTL for 5°, 10° and 20° bin sizes ($p < 0.05$). Column 3 shows that we capture finer regions for which TEN is less than CTL, as we reduce the bin size from 20° to 5°. Column 2 shows the region where TEN is greater than CTL is found when we use a bin size of 5° but not for larger values of bin size. Using a bin size of 5° for the histogram, allows us to identify finer regions that contribute more to the observed changes in ductal network orientation between TEN versus CTL and CONTRA versus CTL. This finding is consistent with the prediction of the optimal bin size from the Wasserstein distance between the distributions for the angular positions as reported in the main text.

**Table S.1.** Differences in the angular positions of the ducts in TEN and CONTRA compared to CTL. The angle ranges in the different columns are those that were significantly different in TEN and CONTRA compared to CTL for different histogram bin sizes ($p < 0.05$).

| bin size | TEN > CTL | TEN < CTL | CONTRA > CTL | CONTRA < CTL |
|---|---|---|---|---|
| 5 | -15° to -11° | 50° to 54°<br>55° to 59°<br>85° to 90° | 10° to 14°<br>20° to 24°<br>25° to 29° | -75° to -71°<br>-45° to -41°<br>-40° to -36°<br>-35° to -31° |
| 10 | | 50° to 59° | 20° to 29° | -50° to -41°<br>-40° to -31° |
| 20 | | 50° to 69° | 10° to 29° | -50° to -31° |
